# Quantifying glioblastoma drug penetrance from experimental data

**DOI:** 10.1101/650184

**Authors:** Susan Christine Massey, Javier Urcuyo, Bianca Maria Marin, Jann Sarkaria, Kristin R. Swanson

**Affiliations:** Mayo Clinic, Phoenix, AZ 85054 USA; Mayo Clinic, Rochester, MN 55905 USA

## Abstract

Poor clinical trial outcomes for glioblastoma (GBM) can be attributed to multiple possible causes. GBM is heterogeneous, such that there is a chance of treatment–resistant cells coming to predominate the tumor, and due to the blood brain barrier (BBB) it is also possible that therapy was inadequately delivered to the tumor. Mathematically modeling the dynamics of therapeutic response in patient–derived xenografts (PDX) and fitting the mathematical model to bioluminescence imaging flux data, we may be able to assess the degree to which both drug resistance and drug penetrance are driving varied responses to these therapies.

## I. Introduction to the type of problem in cancer

Glioblastoma (GBM) is an aggressive primary brain cancer noted for its diffuse infiltration into surrounding normal–appearing brain. This invasiveness makes GBM notoriously difficult to treat, as diffusely invading cells cannot be resected surgically, are difficult to target with radiation therapy, and thus must be targeted with chemotherapy. However, this too presents a challenge, as these invading GBM cells reside beyond the dense tumor regions where angiogenesis causes disruption of the blood brain barrier (BBB) and allows drugs to more readily enter the central portion of the tumor. Thus, failed trials involving molecularly targeted therapies face a daunting task of understanding whether the root cause was inadequate targeting, resistance, or insufficient delivery across the BBB to the tumor. In order to improve treatment outcomes, it is critical to determine predictors of drug distribution in individual patients’ tumors and surrounding brain tissue to ensure invading GBM cells are adequately exposed to the therapy.

Using GBM patient-derived xenograft (PDX) lines to recapitulate the both the inter- and intratumoral heterogeneity seen within and across patients [1,2], several treatments were administered across subjects implanted with different PDX cell lines (derived from different GBM patients) implanted either in flank or orthotopically. The size of PDXs were determined non-invasively using bioluminescence imaging (BLI) flux, which is directly proportional to cell number. The most promising treatment results for the flank tumors was ABT414 (Depatuxizumab Mafodotin), an investigational EGFR-targeted monoclonal antibody drug conjugate [3]. However, the murine flank and orthotopic EGFRmut PDXs treated with ABT414 showed response differences between the two tumor locations for different PDX lines, suggesting a BBB role. Following these experiments, we compiled the time series BLI data from PDXs to develop and parameterize a mathematical model of the observed treatment response dynamics. Fitting our model to this data via nonlinear regression allows us to obtain parameter estimates that can help assess the degree to which these results might be attributed to either the evolution of therapeutic resistance or differences in blood brain barrier breakdown between the tumors.

## II. Illustrative Application of Methods

### A. Ordinary Differential Equation Model Development

BLI data from untreated groups indicated that tumors grew exponentially in terms of total tumor cell number. In the treated groups, however, there was a decline in BLI flux until there appeared to be a subsequent phase of exponential re-growth, albeit with a slower growth rate. As this appeared to indicate a resistant subpopulation of tumor cells, the minimal differential model of tumor growth includes two tumor cell populations corresponding to those that are sensitive (*s*) and resistant (*r*) to the antibody (*A*), as well as the dynamics of the antibody itself:

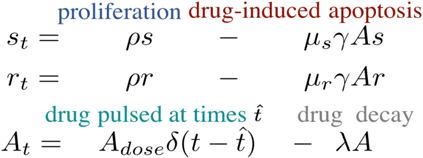

This model is schematized in Fig. 1. Model parameters and their definitions are outlined in Table I.

**Figure 1.**
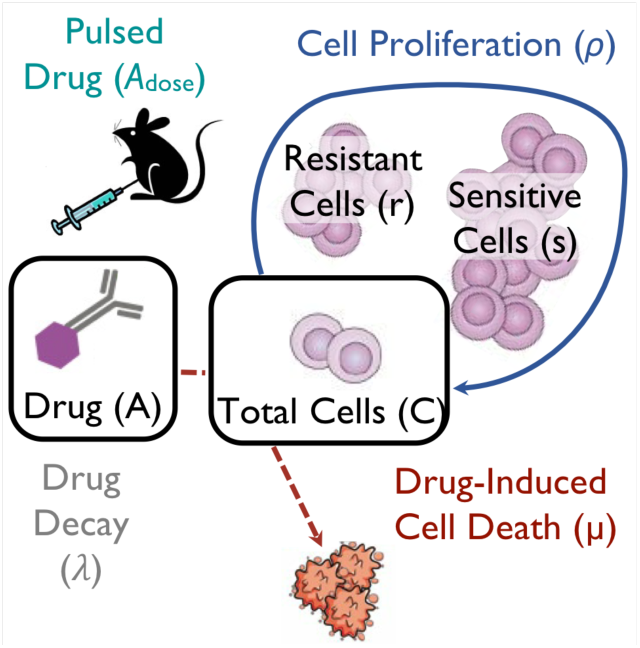
Schematic of patient–derived xenograft response to therapy with ABT414 antibody drug conjugate, including key variables and parameters of the mathematical model.

**TABLE I.** Parameter Symbols and Definitions

| Sym | Parameter Details |  |  |
| --- | --- | --- | --- |
|  | Parameter Definition | Value | Units |
| $\rho$ | cellular proliferation rate | 0.1 to 0.3 | day<sup>-1</sup> |
| $\mu_s$ | drug-mediated sensitive cell kill rate | 2 to 10 | mg<sup>-1</sup> day<sup>-1</sup> |
| $q$ | resistant implanted cell proportion* | 0 to 1 | — |
| $z$ | reduced sensitivity factor in r cells* | 0 to 1 | — |
| $\lambda$ | ABT414 decay rate<sup>a</sup> | $\ln(2)/7$ | day<sup>-1</sup> |
| $A_{dose}$ | ABT414 dose in a single pulse<sup>a</sup> | 0.1 | mg |
| $\gamma$ | Fraction exposed to ABT414<sup>b</sup> | 0 to 1 | — |

a. These parameters were known, all others were found by fitting to data. b. Flank tumor has no blood brain barrier, so $\gamma = 1$ in that setting. (\*Indicates values were constrained to this range for fitting.)

Further, this model can be solved analytically, combining *s* and *r* cells to obtain the total population of tumor cells, *C*, letting *q=r*_*0*_*/C*_*0*_ to represent the proportion of implanted cells thatare resistant, and *z=µr/µ*_*s*_ to represent the degree to which the resistant cells are less sensitive to antibody than the so-called sensitive cells:

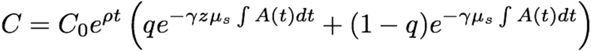

where,

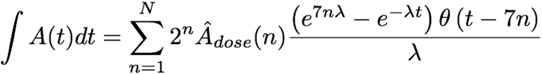

Many parameters were unknown, but could be determined through several steps of fitting the equations to the time series BLI data (as described in II. B.). The only known parameters were the doses of ABT414 delivered, including the timing of dose pulses, as well as the half-life of the drug.

### B. Parameter Estimation by Fitting Model to Data

Starting with the untreated (sham control) case, the treatment components of the model go away, leaving a simple exponential equation *C=C*_*0*_*e*^*ρt*^, which can be fitted to the untreated BLI data to obtain an estimate of viable implanted cells (initial condition *C*_*0*_) and the net growth rate of cells, *ρ* for each PDX line (Fig. 2).

**Figure 2.**
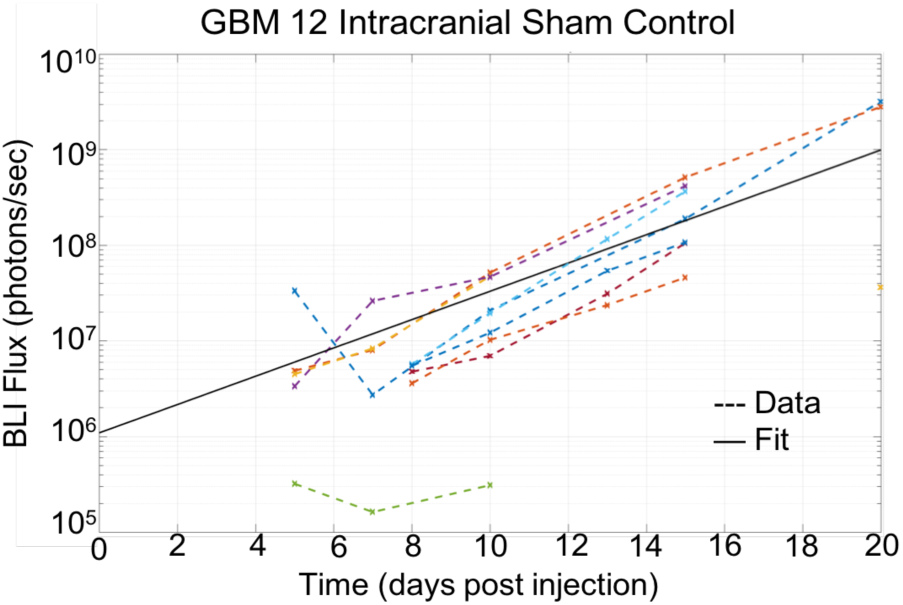
Plot showing untreated tumor growth assessed by bioluminescence imaging flux as well as the untreated model fit found by nonlinear regression.

Next, a similar fitting process was done with treated flank tumors, using these parameter estimates from the untreated case and setting parameter *γ* (representing BBB permeability) to one, since there is no BBB effect to reduce the amount of drug and resultant effect in the tumor. Linear regression performed well (Fig. 3) and found estimates of *q, z* and *µ*_*s*_.

**Figure 3.**
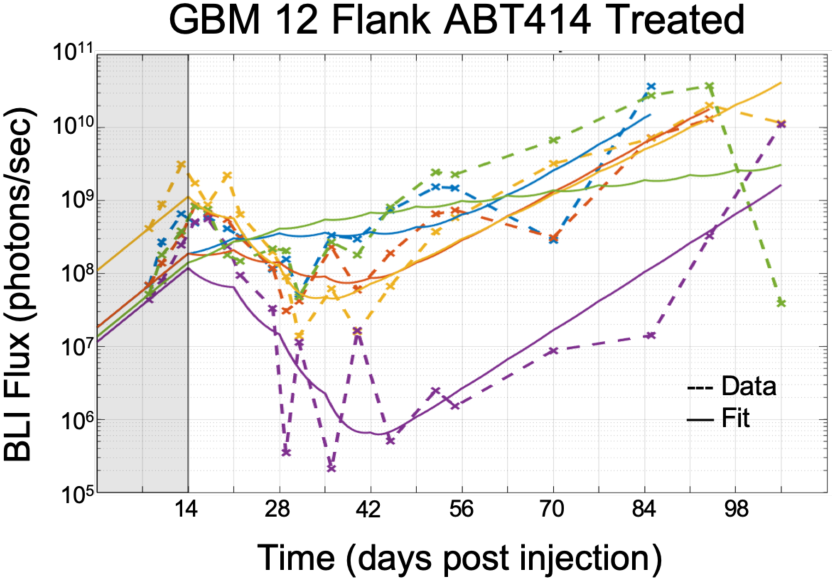
Treated flank tumor growth assessed by bioluminescence imaging flux as well as the individual model fits. Shaded region indicates time before treatment initiated

Finally, keeping *µ*_*s*_ from the flank data (i.e., assuming that a subpopulation remained just as sensitive intracranially as in flank), least squares regression was used to estimate *q, z*, and *γ* (Fig. 4).

**Figure 4.**
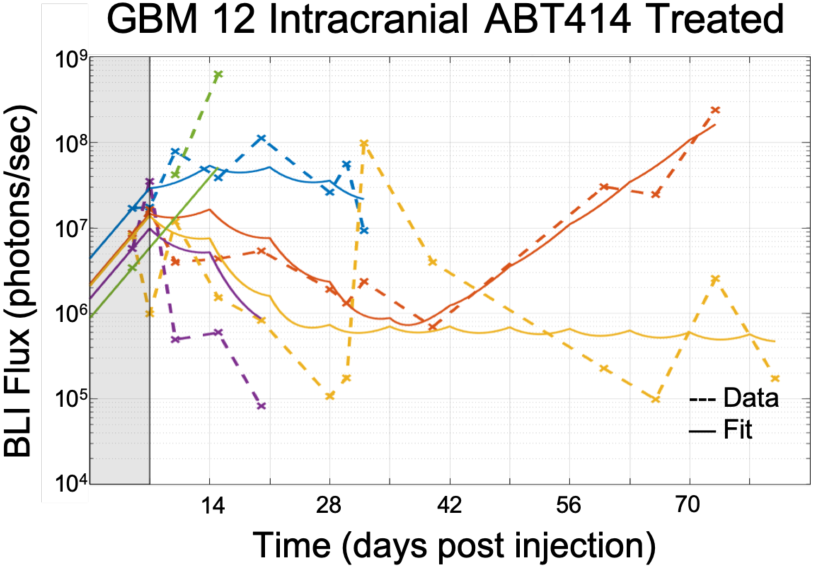
Treated intracranial tumor growth assessed by bioluminescence imaging flux as well as the individual model fits. Shaded region indicates time before treatment initiated.

In spite of the noise in the BLI data, the fits in both the intracranial and flank subjects performed reasonably well. This suggests that we can use this method across all of the PDX lines to quantify differences in sensitivity (including the cell kill rates and proportion of sensitive vs resistant cells initially implanted) and the global fraction of tumor exposed to drug. Ultimately, we anticipate that this modeling framework could be used for assessing the contributions of drug resistance and penetrance in PDXs for other therapeutic agents.

